# EZH2 inhibition sensitizes IDH1R132H mutant gliomas to histone deacetylase inhibitor

**DOI:** 10.1101/2023.07.31.551045

**Authors:** Lisa Sprinzen Houghton, Franklin Godinez Garcia, Angeliki Mela, Liang Lei, Pavan Upadhyayula, Aayushi Mahajan, Nelson Humala, Lisa Manier, Richard Caprioli, Alfredo Quiñones-Hinojosa, Patrizia Casaccia, Peter Canoll

## Abstract

Isocitrate Dehydrogenase-1 (IDH1) is commonly mutated in lower grade diffuse gliomas. The IDH1R132H mutation is an important diagnostic tool for tumor diagnosis and prognosis, however its role in glioma development, and its impact on response to therapy, is not fully understood. We developed a murine model of proneural IDH1R132H mutated glioma that shows elevated production of 2-Hydroxyglutarate (2-HG) and increased tri-methylation of lysine residue K27 on histone H3 (H3K27me3) compared to IDH1 wild-type tumors. We found that using Tazemetostat to inhibit the methyltransferase for H3K27, Enhancer of Zeste 2 (EZH2), reduced H3K27me3 levels and increased acetylation on H3K27. We also found that, although the histone deacetylase inhibitor (HDACi) Panobinostat was less cytotoxic in IDH1R132H mutated cells (either isolated from murine glioma or oligodendrocyte progenitor cells infected in vitro with a retrovirus expressing IDH1R132H) compared to IDH1-wildtype cells, combination treatment with Tazemetostat is synergistic in both mutant and wildtype models. These findings indicate a novel therapeutic strategy for IDH1-mutated gliomas that targets the specific epigenetic alteration in these tumors.

**Main Points:** Murine gliomas initiated by the IDH1R132H mutation (in the presence of additional genetic alterations, such as p53 loss and PDGF overexpression) recapitulate the metabolic and transcriptional features of the proneural subtype, as they are characterized by increased 2HG levels, and are enriched for OPC lineage-restricted genes compared to IDH-wildtype murine gliomas. In murine IDH1-R132H glioma cells, EZH2 inhibition is not cytotoxic as a monotherapy but reduces levels of H3K27me3 and increases levels of H3K27ac. IDH1R132H cells are relatively resistant to Panobinostat cytotoxicity compared to IDH-wildtype cells, but combining treatment with EZH2 inhibition synergistically kills glioma cells and increases H3K27ac.

## Introduction

Diffuse gliomas are the most common type of primary brain tumor and include several biologically distinct tumor types with different molecular and genetic features. The current WHO classification system for CNS tumors incorporates common genetic alterations, and diffuse gliomas are classified based on the presence or absence of mutations in the metabolic enzyme Isocitrate Dehydrogenase (IDH), the most common being IDH1-R132H^1^. Glioblastoma (GBM), the most common and malignant primary brain tumor, is IDH-wildtype, whereas IDH-mutant gliomas are classified as oligodendroglioma or astrocytoma. The majority of IDH-mutant gliomas have a proneural transcriptional phenotype; the gene expression profiles of these cells resemble oligodendrocyte progenitor cells (OPCs) or neural progenitor cells (NPCs) ^2–4^. As such, OPCs are considered a likely cell of origin for IDH-mutant gliomas^5–8^. While the cell of origin in gliomagenesis is still disputed, the proliferation ability, distribution, and abundance in the adult brain further implicates OPCs in this process ^9–14^. Our lab and others have established the tumorigenic potential of OPCs residing in the adult white matter of mice and rats ^5,8,10,15–18^. Although the transforming effects of mutations in IDH have been extensively investigated, the specific effects of mutant IDH1 in cellular transformation of OPCs has not been established. The connection between tumor genotype and phenotype suggests that the transforming effects of IDH-mutation are highly dependent on cellular context, and that OPCs are uniquely susceptible, although the mechanism is not understood.

The ability for OPCs to acquire the expression profile of myelinating oligodendrocytes requires the repression of several transcripts related to other lineages, which is initiated by deacetylation and methylation of specific lysine residues on histone H3 ^19–21^. Enhancer of Zeste 2 (EZH2) is the main writer for H3K27 di- and tri-methylation ^22^ and has a role in OPC lineage specification and proliferation as well as restriction of neuronal and astrocytic lineage commitment ^23^. As such, the intrinsic functional necessity for epigenetic regulation and maintenance of myelination in the adult brain imparts OPCs with an innate susceptibility to metabolic stress.

Wildtype IDH1 produces α-ketoglutarate (α-KG), however mutant IDH1 (IDH1 R132H) produces 2-hydroxyglutarate (2-HG), which acts as a competitive inhibitor of α-KG-dependent demethylases, and in this way IDH1 mutations increase Histone H3 lysine residue 27 trimethylation (H3K27me3) ^24–26^. H3K27 can also be acetylated, however acetylation and trimethylation of the same residue are mutually exclusive ^6,27–31^, including GBM ^32^, and blocking its activity in some human cancer cell lines inhibits growth ^33,34^, suggesting unchecked EZH2 activity causes H3K27 hypermethylation and tumor-suppressor inhibition ^35^. Histone deacetylase inhibitors (HDACi) increase acetylation and have become a promising avenue for cancer treatment ^36–39^. We hypothesized that treating cells with an EZH2 inhibitor (Tazemetostat) and a histone deacetylase inhibitor (Panobinostat) would synergistically increase acetylation to target epigenetic alterations in brain tumors.

## Results

### Retroviral delivery of IDH1R132H mutation with PDGF and p53 deletion induces glioma formation with increased 2HG

To determine the influence of IDH1 mutation on gliomagenesis induced by loss of p53 and overexpression of PDGF, mice with floxed p53 were stereotactically injected to target the glial progenitors of the subcortical white matter with retrovirus expressing PDGFB and Cre-recombinase, alone or in combination with IDH1R132H expressing retrovirus. These two models allow us to compare tumorigenesis and phenotype between wild-type and IDH1R132H OPC transformation, while controlling for other genetic alterations. Tumor growth was monitored by luciferase imaging, but no difference was seen in the growth kinetics. Both IDH1-mutant and wild-type murine gliomas reached end-stage by tumor morbidity with a median survival of 40 days post-injection (Figure 1A), suggesting the IDH1 mutation did not facilitate or inhibit gliomagenesis in this co-delivery model. Tumors collected at morbidity showed large diffusely infiltrating lesions expanding from the site of injection (Figure 1B). The histology showed that both the wild-type and IDH1R132H tumors resembled malignant high-grade human gliomas. The IDH1 mutant tumors recapitulated the proneural glioma while retaining the mutation (Figure 1B and Supplementary Figure 1A). Furthermore, the mutation did not affect the proliferation index of the fully formed gliomas (Supplementary Figure 1B), consistent with the observation that the IDH1R132H and wildtype glioma models have a similar disease progression.

**Figure 1.**
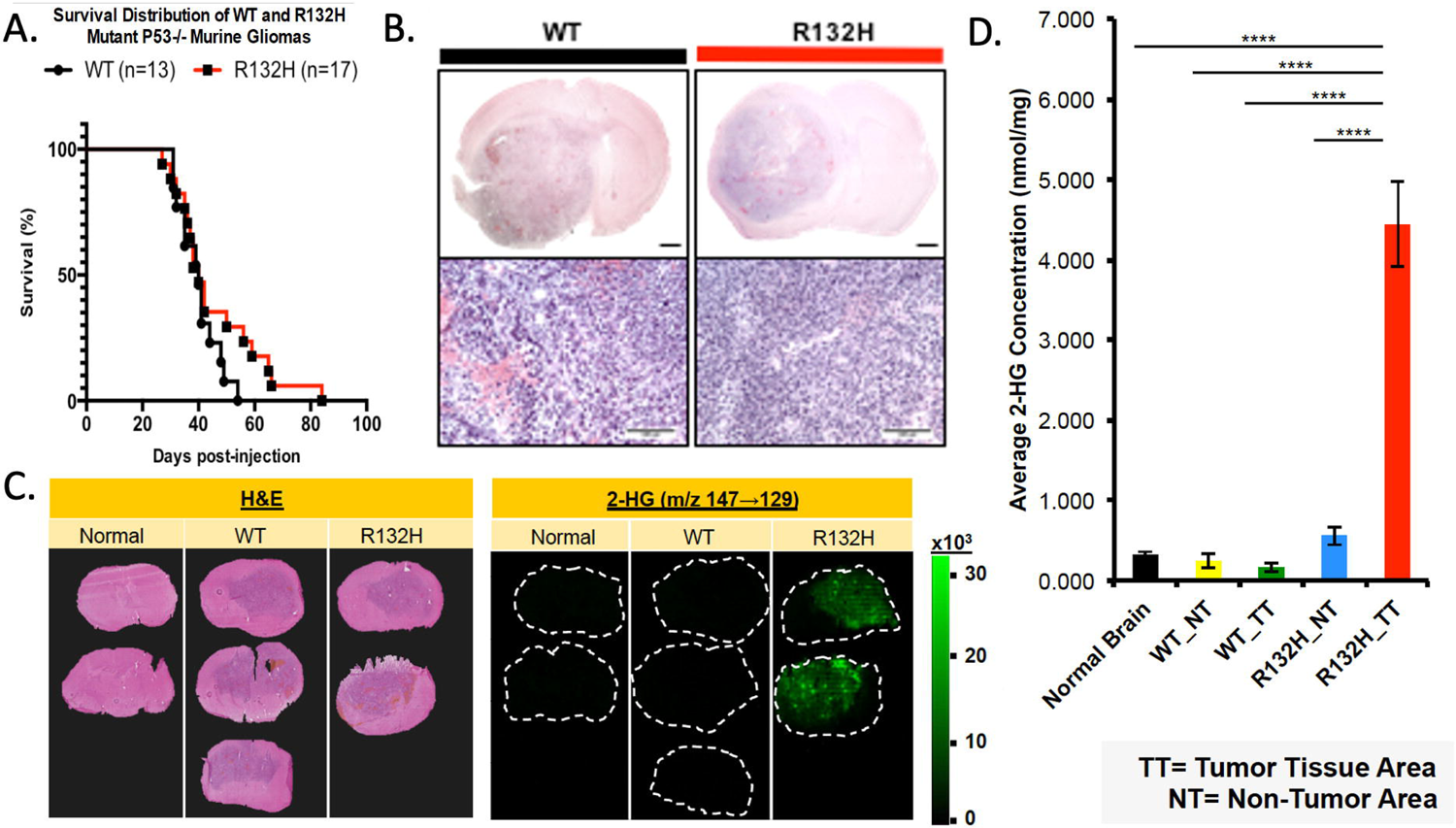
IDH1R132H mutant tumor model does not influence murine survival with increased 2HG. **A)** Kaplan-Meier curve comparing the survival of WT and R132H mouse groups, Log-rank (Mantel-Cox) test p-value = 0.16 and Gehan­ Breslow Wilcoxon test p-value = 0.46. **B)** WT and Rl 32H tumors are histologically indistinguishable by H&E. Upper panel (1.5x) with 1mm scale bar and lower panel (lOx) with lOOµm scale bar. C) Left panel ofH&E staining to show tumor margins. Right panels of MALDI MS two-dimensional ion density images for 2-hydroxyglutaric acid collected at a spatial resolution of 200µm. **D)** Quantification of average concentration (nmol/mg) of 2-HG across tumor (TT) and non-tumor (NT) regions One-way ANOVA with Tukey’s correction for multiple comparison was used.**** p-value <0.0001.

IDH1 normally converts isocitrate to α-KG; however, the R132H mutation changes the enzymatic properties, causing a higher affinity for α-KG rather than isocitrate, thus producing 2HG ^40,41^. Low levels of 2HG are seen under physiological conditions, however IDH1R132H produces elevated levels which have metabolic and epigenetic influences ^42^. We used matrix-assisted laser desorption ionization imaging mass spectroscopy (MALDI MS) to determine if IDH1R132H-expressing tumors had increased 2HG compared to normal brain and wild-type IDH1 tumor tissue^43^. IDH1R132H mutant gliomas showed increased 2HG in the tumor region (Figure 1C/D). We also analyzed α-KG and glutamate levels but found no detectable difference in mutant tumors compared to wild-type or normal brain (Supplementary Figure 1C/D).

### ID1R132H mutation alters the transcriptional phenotype of mouse gliomas

We performed transcriptional profiling by RNA sequencing on the IDH1R132H mutant and wildtype tumors to determine if these tumors would recapitulate the molecular profile of human proneural gliomas. We compared the expression profile of the wildtype (N=5) and IDH1R132H (n=6) murine tumors to the TCGA database using Gene Set Enrichment Analysis (GSEA), with focused analysis on the Verhaak subtype classifier genes: Proneural (PN), Classical (CL), Mesenchymal (MES), Neural-high (NA) and Neural-low (NL) ^2,5,44^. We found that both the IDH1R132H mutant tumors and the wildtype tumors were highly enriched in the PN molecular signature and depleted in the CL and MES signatures (Figure 2A).

**Figure 2.**
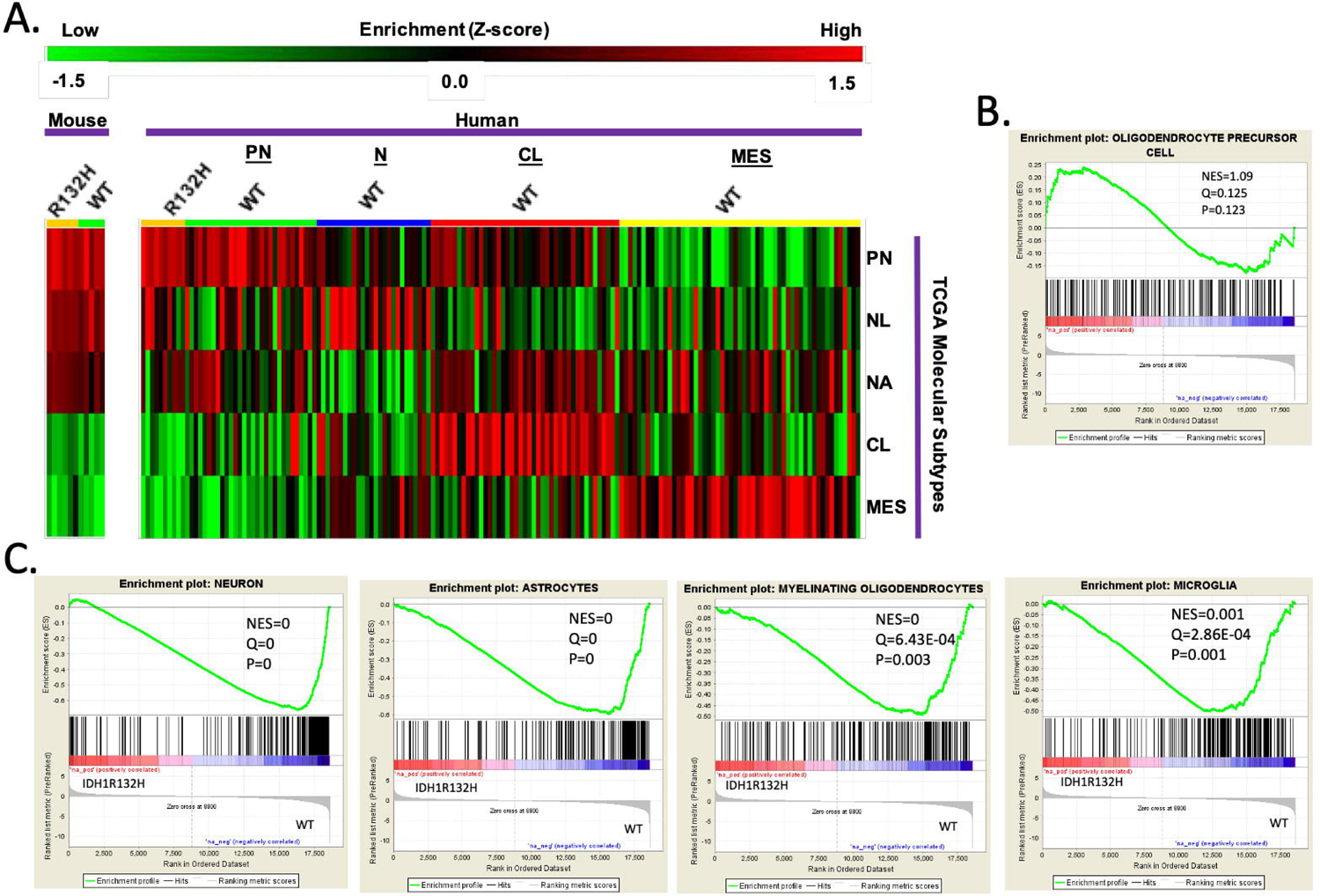
RNA sequencing reveals that wildtype and IDH1R132H murine tumors are proneural and mutant gliomas are more lineage-restricted. **A)** Heatmap of gene-set enrichment analysis (GSEA) used to compare the RNA sequence profiles of IDH1R132H mutant and WT murine gliomas to the four glioma molecular subtypes ^1^ but with splitting Neural into higher or lower expression within the subtype: Proneural (PN), Neural-high (NL), Neural-low (NA), Classical (CL), and Mesenchymal (MES). Samples are grouped as mouse and TCGA human samples. Color scale for high-enrichment (red) and low­ enrichment (green) z-score. **B/C).** GSEA of significant differentially expressed genes using cell-specific brain transcriptome gene sets^2^. Gene sets were made with the top 200 enriched genes based on fold change above average for each cell type: astrocytes, oligodendrocytes, neurons, OPCs, newly formed oligodendrocytes, myelinating oligodendrocytes, microglia, and endothelia. Only showing OPC lineage **(B)** and significant GSEA (neuron, astrocytes, myelinating oligodendrocytes, and microglia) **(C).**

We further analyzed the transcriptional differences between the IDH-mutant and IDH-WT mouse gliomas by performing GSEA using the Barres murine brain transcriptome database ^45^ to create cell-type specific gene sets for astrocytes, oligodendrocytes, neurons, OPCs, newly formed oligodendrocytes, myelinating oligodendrocytes, microglia, and endothelia for analysis. Analysis of the gene sets showed that the IDH1R132H tumors were enriched in OPC genes (Figure 2B), while the IDH-wildtype tumors were significantly enriched in neuronal, astrocytic, myelinating oligodendroglial, and microglial genes (Figure 2C). These findings suggested that the IDH1R132H mutation fostered an OPC-restricted transcriptional phenotype during gliomagenesis, by inhibiting the expression of other neural lineage genes.

### Tazemetostat sensitizes IDH1R132H cells to Panobinostat and combined treatment increases H3K27ac

Based on the increased 2HG levels in the IDH1R132H mutant tumors as well as the transcriptomic data, we wanted to examine if H3K27me3 levels differed between IDH1 mutant and wildtype mouse glioma cells. Western blot analysis of histone extracts from IDH1R132H cells revealed significantly higher H3K27me3 levels compared to IDH1 wildtype cells (Figure 3A/B).

**Figure 3.**
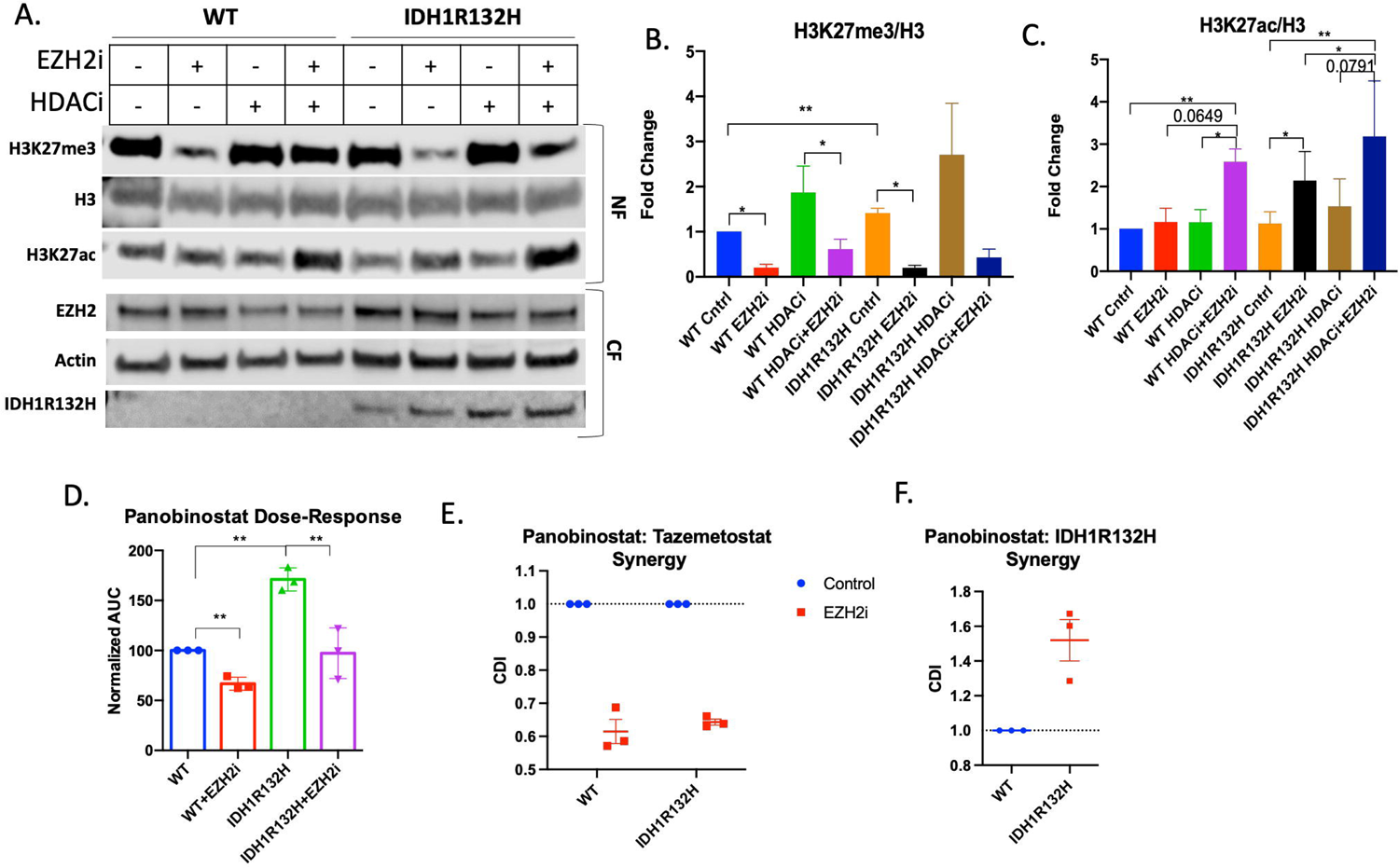
Tazemetostat and Panobinostat co-treatment are synergistic in glioma cells. **A)** Representative western blot of wildtype and IDH1R132H glioma cells were pretreated with 5uM Tazemetostat (EZH2i) or DMSO for 24 hours then retreated with the addition of 20 uM Panobinostat (HDACi) or DMSO for 24 hours. Cells were fractionated for the cytoplasmic fraction (CF) and nuclear fraction (NF) was acid­ extracted. Quantification ofB) H3K27me3 and C) H3K27ac normalized to total H3. All bar graphs are from three independent experiments and shown as fold change to control. Students T-test used for statistical analysis. **D)** Area under the curve analysis from three independent viability experiments normalized to wildtype cells. Paired T-test performed for statistical analysis. **E)** Coefficient of drug interaction from three independent experiments looking at Panobinostat and Tazemetostat synergy in wildtype and IDH1Rl32H cells. **F)** Coefficient of drug interaction from three independent experiments looking at Panobinostat and IDH1R132H.

Based on the differences in post-translational histone modifications, we also compared the viability of our cells after treatment with epigenome-modifying drugs. We found that the wildtype and IDH1R132H mouse glioma cells were equally insensitive to treatment after 72 hours with EZH2 inhibitor Tazemetostat and did not reach IC50 even at 64μM (Supplementary Figure 1E), although treatment was effective in reducing H3K27me3 at doses as low as 0.03μM (Supplementary Figure 1F).

Notably, Tazemetostat (5 μM) significantly reduced H3K27me3 in both IDH1R132H and wildtype cells by 48 hours (Figure 3A/B) and significantly increased H3K27ac in IDH1R132H cells by 48 hours, and in wildtype cells by 116 hours (Supplementary Figure 1G), revealing that Tazemetostat has reciprocal effects on methylation and acetylation at H3K27 in both cell lines.

We used a histone-deacetylase inhibitor, Panobinostat (20 μM for 24 hours), to determine if co-treatment with Tazemetostat would synergistically increase H3K27ac. We acutely treated glioma cells with low drug doses and assessed the presence of post-translational modifications (Figure 3A). Most notably, co-treatment in IDH1WT and IDH1R132H glioma cells revealed significantly higher levels of H3K27ac compared to vehicle-treated control cells (Figure 3C). We further demonstrated the synergistic effects of combined treatment on the H3K27 marks via flow cytometry by comparing across treatment conditions the numbers of cells above a specified immunofluorescence threshold for each histone mark (Supplementary Figure 2C/D).

Based on these findings we asked whether combination treatment with Tazemetostat and Panobinostat would be more effective to induce cytotoxicity in glioma cells. Based on a 72-hour viability assay, the IDH1R132H (IC50=47.64) glioma cells were less sensitive to Panobinostat treatment than wildtype (IC50=20.04) glioma cells. However, 24-hour pre-treatment with Tazemetostat followed by co-treatment increased the Panobinostat sensitivity of IDH1R132H cells to similar levels seen with wildtype cells (WT IC50=17.65 and IDH1R132H IC50=22.95) (Figure 3D and Supplementary Figure 2A/B). We used the coefficient of drug interaction (CDI) to determine the nature of drug coaction, in which a CDI value of <1 indicates synergy and a CDI >1 indicates antagonism. We determined that IDH1R132H mutation and Panobinostat are antagonistic (CDI= 1.52) (Figure 3F). These data show that there is synergy between Tazemetostat and Panobinostat in both cell lines (WT CDI=0.61 and IDH1R132H CDI=0.64) (Figure 3E). The glioma cells harboring the IDH1R132H mutation were resistant to Panobinostat but this is rectified by EZH2 inhibiton. This suggests that the synergistic mechanism of action of the combined treatment is to block methylation, allowing for more acetylation than either single treatment alone.

### Combined treatment in OPCs recapitulate the metabolic and epigenetic alterations found in IDH1-mutated gliomas

Given that the IDH1R132H mutation is observed at early stages of glioma formation, we hypothesize that this mutation and the metabolic alterations that it causes have their primary effect on the glial progenitor cells that give rise to gliomas. To test this hypothesis OPCs were isolated from mcherry-luciferase^stop-flox^ p53^flox/flox^ mice and then infected in vitro with retroviruses expressing cre alone (X-cre) to delete p53 or cre and IDH1R132H (IDH1R132H-cre).

To examine the dynamics of H3K27 regulation at early stages of gliomagenesis, we assessed the epigenetic changes that EZH2i and HDACi combination treatment had on early passage (P<6) p53-deleted OPCs (Figure 4A). IDH1R132H-cre OPCs had higher baseline trimethylation than wildtype X-cre OPCs, similar to what we observed in glioma cells derived from our fully-formed IDH1R132H murine tumor model. Tazemetostat monotherapy decreased trimethylation for both retroviral cell conditions (Figure 4B). Both IDH1R132H-cre and X-cre OPCs also had a marked increase in H3K27ac after Tazemetostat and Panobinostat co-treatment, with a 66.2- and 55-fold change for wildtype and IDH1R132H, respectively (Figure 4C). We further demonstrated these findings using flow cytometry (Supplementary Figure 3C/D).

**Figure 4.**
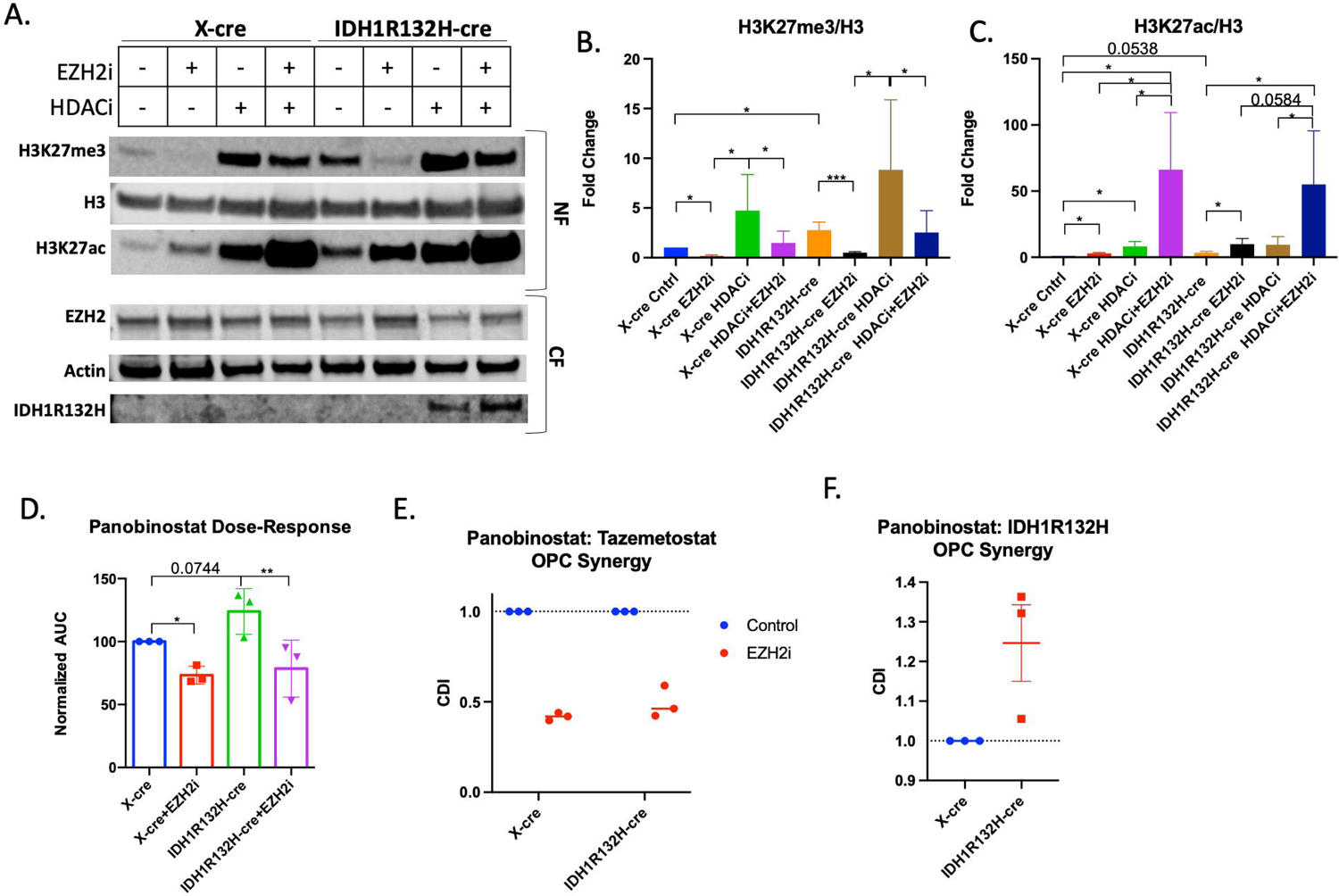
Tazemetostat and Panobinostat co-treatment are synergistic in OPCs. **A)** Representative western blot ofX-cre and IDH1R132H-cre OPCs were pretreated with 5uM Tazemetostat (EZH2i) or DMSO for 24 hours then retreated with the addition of 20 uM Panobinostat (HDACi) or DMSO for 24 hours. Cells were fractionated for the cytoplasmic fraction (CF) and nuclear fraction (NF) was acid-extracted. Quantification ofB) H3K27me3 C) and H3K27ac normalized to total H3. **D)** Area under the curve analysis from three independent viability experiments normalized to wildtype cells. Paired T-test performed for statistical analysis. **E)** Coefficient of drug interaction from three independent experiments looking at Tazemetostat and Panobinostat synergy in X-cre and IDH1R132H-cre infected OPCs. **F)** Coefficient of drug interaction from three independent experiments looking at Panobinostat and IDH1R132H. All bar graphs are from three independent experiments and shown as fold change to control. Students T-test used for statistical analysis.

To determine if Tazemetostat and Panobinostat have a synergistic effect on retrovirally-transformed OPCs, we repeated the combination treatment survival study in wildtype- and IDH1R132H-mutated p53-deleted OPCs. The IDH1R132H-mutated OPCs were less sensitive to Panobinostat (Figure 4D and Supplementary Figure 3A/B), further suggesting mutated IDH1 protects cells from Panobinostat-triggered cell death, despite the higher levels of histone acetylation (WT IC50=15.72 vs IDH1R132H IC50=23.65). Wildtype and IDH1R132H mutant OPCs were more sensitive to the co-treatment than Panobinostat alone (Figure 4D).. Both the wildtype (CDI=0.42) and IDH1 mutant (CDI=0.49) OPCs showed synergy between Tazemetostat and Panobinostat treatment (Figure 4E). We also found that IDH1R132H is antagonistic to Panobinostat treatment (CDI=1.3) (Figure 4F), further suggesting that IDH1R132H protects against the cytotoxic effects of Panobinostat in OPCs. This data demonstrates that the IDH1R132H mutation makes retrovirally-transformed OPCs more resistant to Panobinostat but that combined treatment with Tazemetostat sensitizes cells to Panobinostat.

Together, these results demonstrate that Tazemetostat and Panobinostat co-treatment is synergistic at increasing H3K27ac and reducing cell viability more than either monotherapy in glioma cells and OPCs. Furthermore, the similarities between OPCs and glioma cells suggest that the epigenetic vulnerabilities are inherited from the cell of origin.

## Discussion

In studies that examine the tumorigenic potential of cell-types of the adult brain, OPCs of the adult white matter are seldom explored as candidates that can be selectively susceptible to the formation of gliomas that incorporate an initiating IDH1R132H mutation ^4,46,47^. As such, in this study we designed an approach to compare the epigenetic and transcriptional phenotype of IDH1R132H and IDH1 wildtype glioma model and how early stages of OPC gliomagenesis may provide insight into mechanisms of glial lineage specification that can be therapeutically exploited. We find that while IDH1WT and IDH1R132H tumors had comparable growth dynamics, IDH1R132H recapitulated superphysiological levels of intratumoral alpha-2 hydroxyglutarate and acquired a different transcriptional phenotype characterized by similar levels of OPC-related genes and significantly lower levels of expression for genes associated with neurons, astrocytes, oligodendroglia and microglia. This suggests that IDH1R132H mutation and the associated metabolic and epigenetic alterations restricted cell transformation to an OPC-like phenotype during gliomagenesis.

Although Tazemetostat monotherapy treatment was not cytotoxic in our murine glioma model, it decreased H3K27me3 and increased H3K27ac, which are histone modifications necessary for OPC lineage restriction ^48^. To exploit the cross-talk between acetylation and trimethylation on H3K27, we treated tumor-derived cells with both Tazemetostat and the pan-HDAC inhibitor Panobinostat. We hypothesized that combining an HDAC inhibitor and an EZH2 inhibitor would be synergistic by targeting epigenetic mechanisms (i.e., histone deacetylase and histone methytransferase), whereby EZH2 inhibitors remove H3K27me3 to allow for acetylation of H3K27, which could be beneficial to proneural glioma patient prognosis ^49^. Pharmacologically, we can take advantage of this secondary reciprocal result of EZH2 inhibition and exploit it using an HDAC inhibitor. Others have found that the combination of EZH2 inhibition with HDACi is synergistic in other types of cancer^50–53^, however the effect has previously not been tested in IDH1R132H glioma. Both tumor-derived glioma cells and transformed OPCs show similar responses to EZH2 and HDAC inhibition, suggesting that this mechanism is retained throughout the process of OPC gliomagenesis. Using both Tazemetostat and Panobinostat increased H3K27ac more than either monotherapy and co-treatment was more cytotoxic to the cells than monotherapy specifically in IDH1 mutated cells.

We found that IDH1R132H in combination with other genetic alterations transformed OPCs to proneural glioma. Detailed understanding of epigenetic alterations provides a therapeutic target to leverage. Understanding the cell of origin and how genetic alterations play out within that cellular context, can help understand therapeutic effects and develop new therapeutic strategies. We found that IDH1R132H mutant murine tumors had increased H3K27me3 which rendered those cells resistant to Panobinostat treatment. Treating cells with the EZH2 inhibitor removed H3K27me3 to allow for Panobinostat to be more effective. It is important to note that our model retains both endogenous copies of wildtype IDH1 with IDH1 mutant overexpression, which we acknowledge does not recapitulate physiological gene ratios in human IDH1 mutant tumors. We also looked at global epigenetic changes and not the transcriptional outcome of these changes. Future studies should determine the transcriptional effects of using these drugs in the heterozygous physiological setting. These results show that although an epigenetic modifying drug might not be effective as a monotherapy^54^, it can still be beneficial in combination treatments for glioma. Epigenetic modifying drugs can be more effective in combination by targeting antagonistic modifications (such as methylation and acetylation).

## Methods

### Retrovirus construction

PDGF-IRES-Cre was generated by cloning human PDGF-B and Cre into pQXIX vector (Clontech) as previously described^5^. IDH1R132H-IRES-CRE and PDGF-IRES-GFP retroviruses were generated by cloning the human IDH1R132H sequence and Cre or PDGF-B and GFP into the same vector. VSVG pseudotyped retrovirus was generated as previously described^5^. Briefly, 293GP2 cells were incubated with plasmids expressing VSVG and each viral plasmid mixed with CaCl_2_ and Hepes Buffered saline. Retroviruses were harvested and filtered then re-suspended in PBS.

### Intracerebral stereotactic injections

Brain surgery procedures were done in adult mixed background transgenic mice harboring floxed Trp53 and stop-floxed mCherry Luciferase (Thomas Ludwig, unpublished). For subcortical white matter targeting, we used the coordinates of 2.0 mm lateral, 2.0 mm rostral and 2.0 mm deep, with bregma as the reference point. Using a stereotaxis platform, 1μl of the specified retrovirus was injected using a Hamilton syringe (flow rate 0.2μl/min) at a depth of 2mm. For serial transplantation experiments, 50×10^3^ primary tumor cells were resuspended in Opti-MEM (Invitrogen) and injected at the same coordinate. All injections were done in mice between six and eight weeks of age. Overexpression of the PDGF-BB mitogen, combined with the deletion of the tumor suppressor gene forms brain tumors with the histopathologic and molecular features of proneural glioblastomas. Tumor progression was monitored using Xenogen IVIS Spectrum imager 10-15 minutes after 100ul intraperitoneal injection of 30mg/ml luciferin (Caliper Life Sciences) with 1-minute exposure.

### MALDI Imaging Mass Spectrometry by MS/MS

At tumor end-stage, the whole brain of mice with IDH1R132H and IDH1WT gliomas conditions were harvested and coronally bisected at the retroviral infusion site and posterior to the tumor. The anterior section was used for H&E and IDH1R132H immunohistochemical staining. The coronal tumor-bearing slab was sliced at 12microns and used for acquiring two-dimensional ion density images for glutamic acid (m/z 146 -->128) and 2-hydroxyglutaric acid (m/z 147-129) on a MALDI LTQ XL linear ion trap mass spectrometer at a spatial resolution of 50microns according to methods previously described^43^. Solution standards of 2-hydroxyglutaric acid concentrations were used for generating a standard calibration curve (intensity ratio of 2-HG/ internal standard vs concentration of 2-HG (pmol/μL) using weighted linear regression (1/X^2^).

### Primary OPC cultures

Primary mouse OPCs were isolated from the brain of floxed p53, stop-floxed mCherry Luciferase mice at postnatal day 5 through immunopanning with a rat anti-mouse CD140a antibody, recognizing PDGFRα, as previously described by Dr. Patrizia Casaccia’s laboratory^55,56^, and were cultured in a modified SATO medium (Dulbecco’s modified Eagle’s medium (DMEM), 4 μg/ml selenium, 5 mg/ml insulin, 1mM sodium pyruvate, 2mM L-glutamine, 100 U/ml penicillin, 100 g/ml streptomycin, B27 Supplement, N2 supplement, Trace Element B, 10 μg/ml biotin) supplemented with PDGF-AA (20 ng/ml) and basic fibroblast growth factor (bFGF, 20 ng/ml). Trp53fl/fl OPCs were cultured as described above and then infected with an X-IRES-CRE or IDH1R132H-IRES-CRE retrovirus to obtain Trp53−/− OPCs and IDH1R132H Trp53−/− OPCs respectively.

### Primary tumor cell lines

Primary tumor cell lines were derived as described previously^5,57,58^. Once mice showed evidence of neurological features consistent with tumor morbidity, mice were sacrificed by cervical dislocation and decapitation. Brains were acutely harvested and maintained in prewarmed PBS (37°C). Tumors were surgically resected, minced in 2.5% Trypsin and serially triturated by syringe using 18G and 21G needles. Triturated tumor tissue was filtered (70µm) then neutralized in 50% FBS and centrifuged at 1500rpm for 5 minutes. Supernatant was discarded and tissue was cultured in defined tissue culture media containing high glucose DMEM (5.4g/L D-glucose, L-glutamine), 0.5% FBS, N2 Supplement, PDGF-AA (10 ng/ml), bFGF (10 ng/ml), and antibiotic–antimycotic solution.

### Drug Treatment Cell Viability Assay

Cells were plated in black-walled 96-well plates and allowed to settle for 24 hours. For combined treatment studies: cells were pre-treated with Tazemetostat (5uM) or DMSO for 24 hours and treated by doubling serial Panobinostat dilutions (1-128nM) with Tazemetostat or DMSO for 72 hours. CellTiter-Glo (Promega) was added to cells following manufacturer’s protocol. The luminescence was read on GloMax microplate reader (Promega) using provided manufacturer’s protocol.

### Drug Combination Treatments

For Western Blot and Flow Cytometry studies, cells were plated and allowed to settle for 24 hours. Cells were pre-treated with Tazemetostat (5μM) or DMSO for 24 hours. Cells were then co-treated with the addition of Panobinostat (20μM) or DMSO for an additional 24 hours providing four groups: Control (DMSO alone), Tazemetostat (and DMSO), Panobinostat (and DMSO), and Tazemetostat and Panobinostat.

### Whole cell lysis

Cells were washed with ice-cold PBS then scraped in Cell Extraction Buffer (Invitrogen). Cells were moved to Eppendorf tubes and kept on ice for lysis. Tubes were sonicated every 10 minutes for 30 minutes. Samples were spun at 13,000 rpm for 10 minutes at 4 °C and the lysate was transferred to new tubes and stored at - 80 °C until use.

### Subcellular protein fractionation from cultured cells

Cells were lysed in hypotonic buffer (10 mM HEPES, pH 7.9, 1.5 mM MgCl_2_,10 mM KCl) supplemented with 0.5 mM dithiothreitol (DTT), 1 mM phenylmethylsulfonyl fluoride (PMSF), 5mM sodium butyrate, phosphatase inhibitor cocktail, and protease inhibitor cocktail freshly prepared (at 4 °C for 15 min), followed by 0.5% NP40 treatment (vortex for 10 s) to disrupt the cell membranes. Lysates were then centrifuged at 1500 × g for 10 min at 4 °C to separate the cytoplasmic components (supernatant) from the nuclei-enriched fractions (pellets). Volumes (0.11μL/μL) of 10× cytoplasmic extraction buffer (0.3 M HEPES, 1.4 M KCl, and 30mM MgCl_2_) was added to the supernatant and sonicated for 30 s ON/OFF for 5 min at high power in Bioruptor (Diagenode). After centrifugation at 16,000 × g for 10 min at 4 °C, the soluble fraction was collected as a cytoplasmic extract, and the pellet as the nuclear fraction.

### Differential centrifugation subcellular protein fractionation from brain homogenate

Tissue samples were homogenized in 50mM HEPES, 125mM NaCl, 0.1M sucrose with fresh protease inhibitor cocktail. Samples were spun for 10 minutes at 1000 x g at 4 °C and the supernatant was collected as the cytoplasmic fraction and the pellet as the nuclear fraction.

### Histone extraction and purification after fractionation

Histones were extracted from the nuclear fraction using the acid extraction method with all steps performed at 4 °C ^59^. The nuclear pellet was incubated for 1.5 hours to overnight in 0.4 N H_2_SO_4_. After centrifugation at 16,000 x g for 10 min, nuclear debris were removed, and acid-soluble histones were then precipitated using trichloroacetic acid for at least 30 minutes. Samples were spun for 10 minutes 16,000 x g and washed twice in acetone then re-suspended in water.

### Immunofluorescence and immunohistochemistry

Cells for immunofluorescence were washed with PBS then fixed with 4% paraformaldehyde (PFA) for 10 min at room temperature and then the membrane was permeabilized and blocked with 0.5% (vol/vol) Triton X-100 (Fisher Scientific) and 5% normal goat serum for 1 hour. Primary antibodies were applied overnight at 4 °C followed by incubation of appropriate secondary antibodies conjugated with fluorophores. Images were captured using Nikon TE2000 epifluorescence microscope equipped with MetaMorph software (Version. V.7.6.5.0, Molecular Devices). Quantification of the immunofluorescent intensity was done using ImageJ. Tumor bearing mice were anesthetized and perfused with 4% PFA. After tissue processing and paraffin embedding, sections of 5 μm were cut sectioned by the Columbia University Medical Center Molecular Pathology Shared Resource facility. To perform immunohistochemistry, sections were de-paraffinized, immersed in 10 mM citrate buffer, pH 6.0, for 10 min in the microwave at 650 W, followed by blocking with 10% normal goat serum, before overnight incubation of primary antibodies at 4 °C. Appropriate secondary antibodies conjugated with fluorophores were used the following day to complete the staining. DAPI (4′,6-diamidino-2phenylindole) was used as a nuclear counterstain.

### Microscopy

Stained tissue sections and fluorescent reporters of labeled tumor-derived cells were imaged using a Nikon TE2000 epifluorescence microscope equipped with MetaMorph software (Version. V.7.6.5.0, Molecular Devices). Micrographs were processed and merged using MetaMorph and ImageJ.

### Cellular Dissociation for Flow Cytometry

Slice culture and brain tumors were dissociated using filtered Carica Papain extract diluted 1:20 in 1N NaOH PBS (supplemented with 2mg/mL L-cysteine) for 30 minutes at 37°C with shaking then centrifuged in excess PBS at 400xg for 5 minutes. The slices were manually dissociated with a pipette then sucrose was added (final 15%) and the samples were spun at 1000 rpm for 5 minutes to remove debris. The remaining bottom ∼30% was washed in PBS then processed for flow cytometry.

### Flow cytometry

Cells were collected and fixed with 4% formaldehyde (methanol-free) for 15 minutes at room temperature. Cells were centrifuged in excess PBS 400 x g for 5 minutes for all washes. Cells were permeabilized by adding 100% ice-cold methanol drop-wise while gently vortexing cells to a final concentration of 90% methanol. Cells were incubated on ice for 30 minutes then washed in excess PBS to remove methanol. Cells were resuspended in diluted primary antibodies and incubated for 1 hour at room temperature. They were washed in excess PBS and then resuspended in diluted fluorochrome-conjugated secondary antibodies and incubated for 30 minutes at room temperature. Cells were washed again in excess PBS then resuspended in PBS for flow cytometry reading. The BD LSRFortessa was used for optical measurements and analysis was performed by FlowJo version 9 (Ashland, OR: Becton, Dickinson and Company; 2019).

### RNA extraction and sequencing

Total mRNA was extracted using the Qiagen RNeasy Mini kit (Qiagen, 74106). The Columbia Sulzberger Genome Center sequenced the RNA samples (mm10, Illumina iGenomes) and Tophat2 was used to map the reads. Differential expression analysis was performed using Cuffdiff2. We used expression data from the murine brain transcriptome database^45^ to create cell type specific gene sets with the top 200 enriched genes, based on fold change above average expression, found in astrocytes, oligodendrocytes, neurons, OPCs, newly formed oligodendrocytes, myelinating oligodendrocytes, microglia, and endothelia. We performed Gene Set Enrichment Analysis^44^ on the RNA sequencing data using on these cell type specific gene sets.

### Statistical Analysis and Graphing

The Coefficient of Drug Interaction (CDI) was used to determine if drug:drug and drug:mutation interactions were synergistic (CDI<1) or antagonistic (CDI>1)^60,61^. The CDI was calculated using CDI=AB/(A x B), where AB is the OD value ratio of the combination group to control group and A or B is the OD value ratio of a single agent to control group. All graphs with statistical analysis were made in Graphpad Prism version 8.0.0 for Mac, GraphPad Software, San Diego, California USA, www.graphpad.com. All volcano plots and statistical analysis were made using R Core Team (2020). R: A language and environment for statistical computing. R Foundation for Statistical Computing, Vienna, Austria, www.R-project.org. Heatmaps were made using R or Multi-experiment viewer (MeV) version 10.2 ^62^. Western blots were analyzed and quantified using Image Studio Lite analysis software V. 5.2 (LI-COR Biosciences).

## Supporting information

Mouse glioma GSEA

Mouse GSEA TCGA

Materials and Methods - Antibodies

---

**Supplementary Figure 1:**
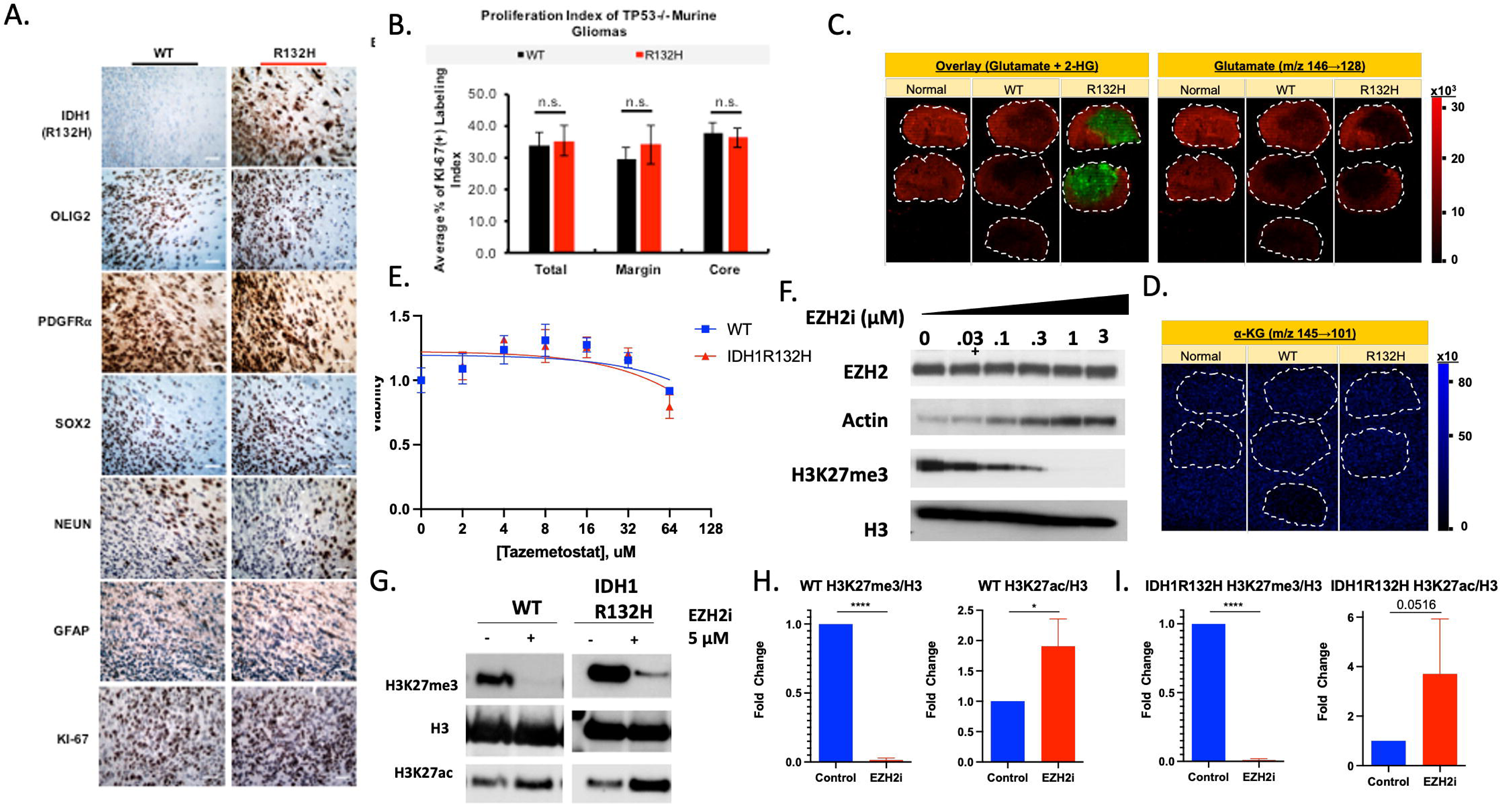
**A)** Panel of micrographs comparing immunohistology staining of WT and RI 32H murine gliomas for markers of proneural glioma phenotype (OLIG2 and PDGFRa), R132H mutation, neuronal marker (NEUN), astrocyte lineage marker (GFAP), proliferation marker (Ki-67), and tumor region (Sox2). Scale bars correspond to 50µM at 20X magnification. **B)** Ki-67 labeling indexes of end-stage wildtype and IDH1R132H mutated gliomas at the core (N=3), margin (N=3), and total. Quantification was performed by counting,_,1,000 cells per tumor experimental condition as a percent ofki-67 positive cells. MALDI-IMS two-dimensional ion density images of normal mouse tissue, WT and IDH1R132H murine gliomas for C) Glutamate (right panel) and Glutamate 2-hydroxyglutarate overlay (left) and **D)** a-ketoglutaric acid. E). Dose vs. response of wildtype and IDH1R132H mutation glioma cells with increasing doses of Tazemetostat (Log2 scale). Viability determined by Cell Titer Glo assay normalized to untreated control. Line represents nonlinear fit. **F).** Western blot of whole cell lysate showing decreasing levels ofH3K27me3 with increasing doses ofTazemetostat after 72 hours in wildtype glioma cells. **G).** Representative western blot of wildtype and IDH1R132H glioma cells after 116 hours ofTazemetostat (5µM) and histone acid extraction. Quantification of wildtype **H)** and IDH1R132H I) glioma cell H3K27me3 and H3K27ac levels normalized to total H3 from three independent experiments. Represented as fold change to control for each experiment. Error bars correspond to SEM and Student’s t-test used for statistical analysis (n.s: p=value>0.05).

**Supplementary Figure 2:**
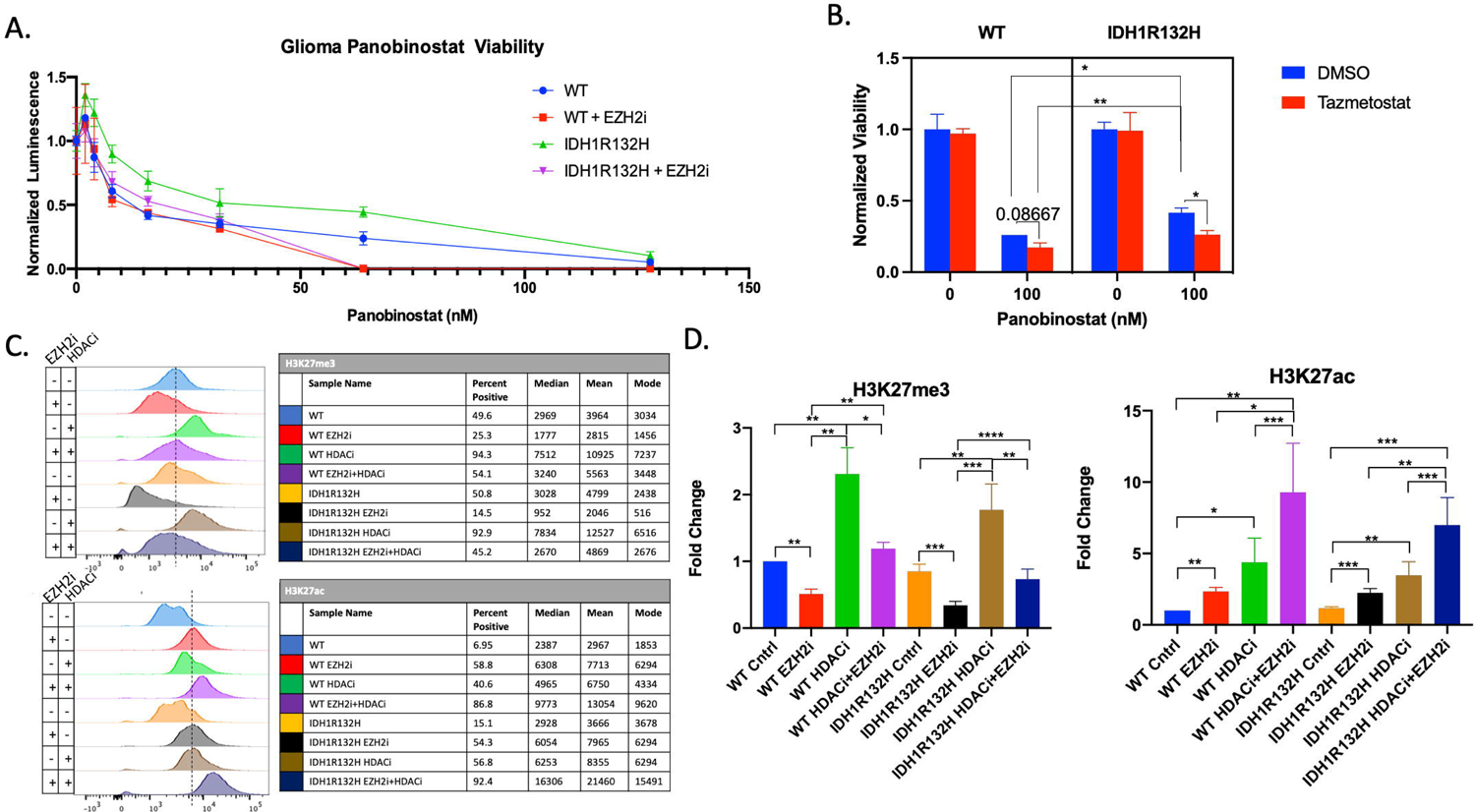
**A)** Representative dose-response ofwildtype and 1D1R132H glioma cells treated for 72 hours with serial dilutions of Panobinostat with or without 5uM Tazemetostat (EZH2i). Luminescence values normalized to untreated cells from Cell Titer Glo. **B)** Representative 3-way ANOVA ofwildtype and IDH1R132H glioma cells with multiple comparisons. Cells were plated and pre-treated with DMSO or Tazemetostat and then treated with DMSO or lOOnM Panobinostat for 24 hours. Source of variation: Panobinostat (94.94% of variation, P<0.0001), IDH1R132H (0.8331% of variation, P=0.0234), Tazemetostat (0.9185% of variation, P=0.0182), Panobinostat x Tazemetostat (0.6019% of variation, P=0.0491). **C)** Representative flow cytometry analysis ofH3K.27me3 and H3K.27ac stained glioma cells. Table shows the percent positive cells based on dotted line, the median, mean, and mode. **D)** Quantification from fold change ofH3K.27me3 and H3K.27ac mean. All bar graphs are from three independent experiments and shown as fold change to control. Students T-test used for statistical analysis

**Supplementary Figure 3:**
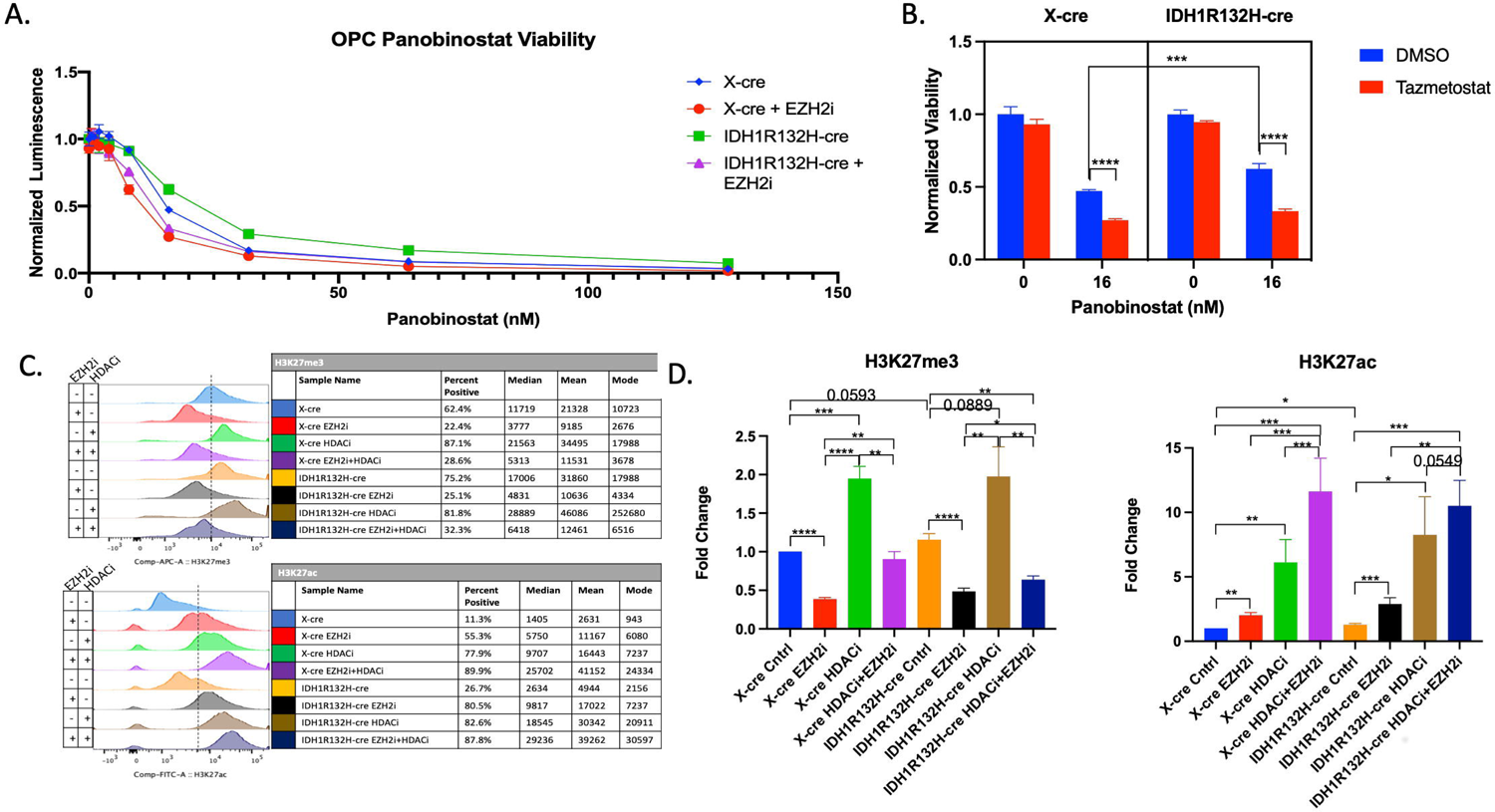
**A)** Representative dose-response of X-cre and ID1R132H-cre infected OPCs treated for 72 hours with serial dilutions of Panobinostat with or without 5uM Tazemetostat (EZH2i). Luminescence values normalized to untreated cells from Cell Titer Glo. **B)** Representative 3-way ANOVA ofwildtype and IDH1R132H glioma cells with multiple comparisons. Cells were plated and pre-treated with DMSO or Tazemetostat and then treated with DMSO or 16nM Panobinostat for 72 hours. Source of variation: Panobinostat (87.77% of variation, P<0.0001), IDH1R132H (0.9731% of variation, P=0.0002), Tazemetostat (7.045% of variation, P<0.0001), Panobinostat x IDH1Rl32H (0.7463% of variation, P=0.0006), Panobinostat x Tazemetostat (2.512% of variation, P<0.0001), Panobinostat x IDH1R132H x Tazemetostat (0.1968% of variation, P=0.0429). C) Representative flow cytometry analysis ofH3K27me3 and H3K27ac stained glioma cells. Table shows the percent positive cells based on dotted line, the median, mean, and mode. **D)** Quantification from fold change ofH3K27me3 and H3K27ac mean. All bar graphs are from three independent experiments and shown as fold change to control. Students T-test used for statistical analysis.

